# A decay effect of the bacterial growth rate associated with genome reduction

**DOI:** 10.1101/308817

**Authors:** Kouhei Tsuchiya, Yang-Yang Cao, Masaomi Kurokawa, Kazuha Ashino, Tetsuya Yomo, Bei-Wen Ying

**Author notes:** Corresponding author: Bei-Wen Ying.

## Abstract

Bacterial growth is an important topic in microbiology and of crucial importance to better understand living cells. Bacterial growth dynamics are quantitatively examined using various methods to determine the physical, chemical or biological features of growing populations. Due to methodological differences, the exponential growth rate, which is a parameter that is representative of growth dynamics, should be differentiated. This study experimentally verified the differentiation in growth rates attributed to different methodologies, and demonstrated that the most popular method, optical turbidity, led to the determination of a lower growth rate in comparison to the methods based on colony formation and ATP abundance, due to a decay effect of reading OD_600_ during a population increase. Accordingly, the logistic model, which is often applied to growth data reading the OD_600_, was revised by introducing a new parameter: the decay rate, to compensate for the lowered estimation in growth rates. The modified logistic model not only presented an improved goodness of fit in comparison to the original model but also led to an intriguing finding of a correlation between genome reduction and the decay rate. The decay effect seemed to be partially attributed to the decrease in cell size accompanied by a population increase and was medium dependent. In summary, the present study provides not only a better theoretical tool for the high-throughput studies on bacterial growth dynamics linking with experimental data using optical turbidity to the theoretical analysis with biological importance, but also a valuable insight for understanding the genome evolution and fitness increase in microbial life.

## Introduction

Bacterial growth dynamics have been intensively studied at both the experimental and theoretical levels due to their broad applications in the food and medical industries and because of the simplicity of the fundamental investigation of determining living systems (Yates and Smotzer, 2007; Peleg and Corradini, 2011; Egli, 2015). Theoretically, it is well-known that bacterial growth commonly follows an S-shaped curve on a two-dimensional plane over time the bacterial population. To describe and/or predict S-shaped growth curves, a number of mathematical models have been proposed, such as the Logistic (Verhulst, 1845; 1847) and Gompertz (Winsor, 1932) models, and revised and expanded repeatedly to better understand the biological process of bacterial growth under varied conditions (Fujikawa and Morozumi, 2005; Kargi, 2009; Koseki and Nonaka, 2012; Alonso et al., 2014; Desmond-Le Quemener and Bouchez, 2014; Hermsen et al., 2015). Although the growth curve of the most representative bacterium, *Escherichia coli*, has been examined since the 1930s (Winsor, 1932), the indescribable complexity of the growth dynamics of *E. coli* remain and are still under investigation with other models to understand the current differential growth dynamics (Swain et al., 2016; Tonner et al., 2017).

Despite the variation in theoretical models, only a few biological parameters are known to impact these models. The growth rate during the exponentially growing phase is the most important parameter and has a high biological impact; this rate represents the genetic and environmental influences on the bacterial growth dynamics (Roessler et al., 2003; Ponciano et al., 2005; Liu et al., 2016). Precise evaluation on the growth rate of a growing bacterial population is highly required. Experimentally, bacterial growth is quantified by varied techniques and is dependent on different principles of chemistry, physics and biology (Harris and Kell, 1985; Madrid and Felice, 2005), such as assays of optical turbidity, colony formation, ATP abundance and particle counts (Lin et al., 2010; Myers et al., 2013; Ziv et al., 2013; Pla et al., 2015). The most commonly used method is the conventional method to measure the optical turbidity of a bacterial culture at a wavelength of 600 nm (OD_600_) due to its easy manipulation. To date, this method seems to be the only tool for assessing high-throughput assays of bacterial growth in microplates (Hall et al., 2014; Kurokawa and Ying, 2017), which are commonly used in systematic studies. Several computational techniques have been developed for use in systematic analyses of bacterial growth dynamics (Verissimo et al., 2013; Hall et al., 2014; Sprouffske and Wagner, 2016). These up-to-date studies are largely based on fitting the widely used logistic model to experimental data acquired from temporal reads of optical turbidity (OD_600_). Since the measurement of OD_600_ detects light transmitted through a cell culture by a spectrophotometer, the changes in absorbance may not always accurately correlate to the changes in the number of living cells. Although the optical turbidity was confirmed to be reliable for estimation of bacterial growth rate, the calibration of the optical measurements was required (Dalgaard et al., 1994).

To address how much difference in growth rates caused by different methods, the growth rates of the *Escherichia coli* cells carrying different genomes were evaluated by different methods, in parallel, in the present study. According to the experimental design, the commonly used growth model of the logistic equation was revised by introducing a new parameter of biological importance, which was differed from the previous study that applying a numeral calibration of the measurements (Dalgaard et al., 1994). The fitting efficiency of the modified logistic model was further evaluated using a large data set composed of hundreds of growth curves. As a consequence, a correlation between genome reduction and the logistic parameters was observed for the first time. This finding not only indicates the high impact of the modified logistic model for studying bacterial growth dynamics but also offers intriguing insights on the correlated changes in the cellular features of the population, *e.g.*, an increase in cell size.

## Results and Discussion

### Differences in growth rates evaluated by three commonly used methods

The wild-type *Escherichia coli* strain W3110 was cultured in minimal medium M63 and was temporally sampled in three different assays to measure the optical turbidity (OD_600_), colony formation (CFU) and ATP abundance (Fig. 1A), which are based on the physical, biological and chemical properties of the growing cell population, respectively. Although these three methods are well established and widely used in experimental and theoretical studies on bacterial growth, it is unclear whether they result in a common conclusion regarding bacterial growth. The three methods of measuring the OD_600_, CFU and ATP represent the density of the biomass, number of living cells and cellular activity, respectively. The growth dynamics represented by the changes in these different parameters must be varied in consequence. To verify this assumption, we performed three assays in parallel on identical cell cultures (Fig. 1A). According to the growth curves generated from the OD_600_, CFU and ATP, the growth rates were calculated by regression toward the time sampling records during the exponential growth phase and as described in the Materials and Methods. Repeated cell cultures (N=3~5) showed significant differentiation of the growth rates estimated by measuring the optical turbidity (OD), colony formation (CFU) and ATP abundance (Fig. 1B). The growth rates estimated by measuring the CFU and ATP were comparable, whereas those from OD_600_ were lower (*P*<0.05), regardless of identical population growth. The results demonstrated that the bacterial growth dynamics, which are represented by changes in optical turbidity, were different from those evaluated by changes in the number of active cells, consistent to the previous report (Dalgaard et al., 1994). How and what caused the differences in growth rates and whether the differences were common under other conditions were further investigated by comparing two representative methods, OD_600_ and CFU.

**Figure 1.**
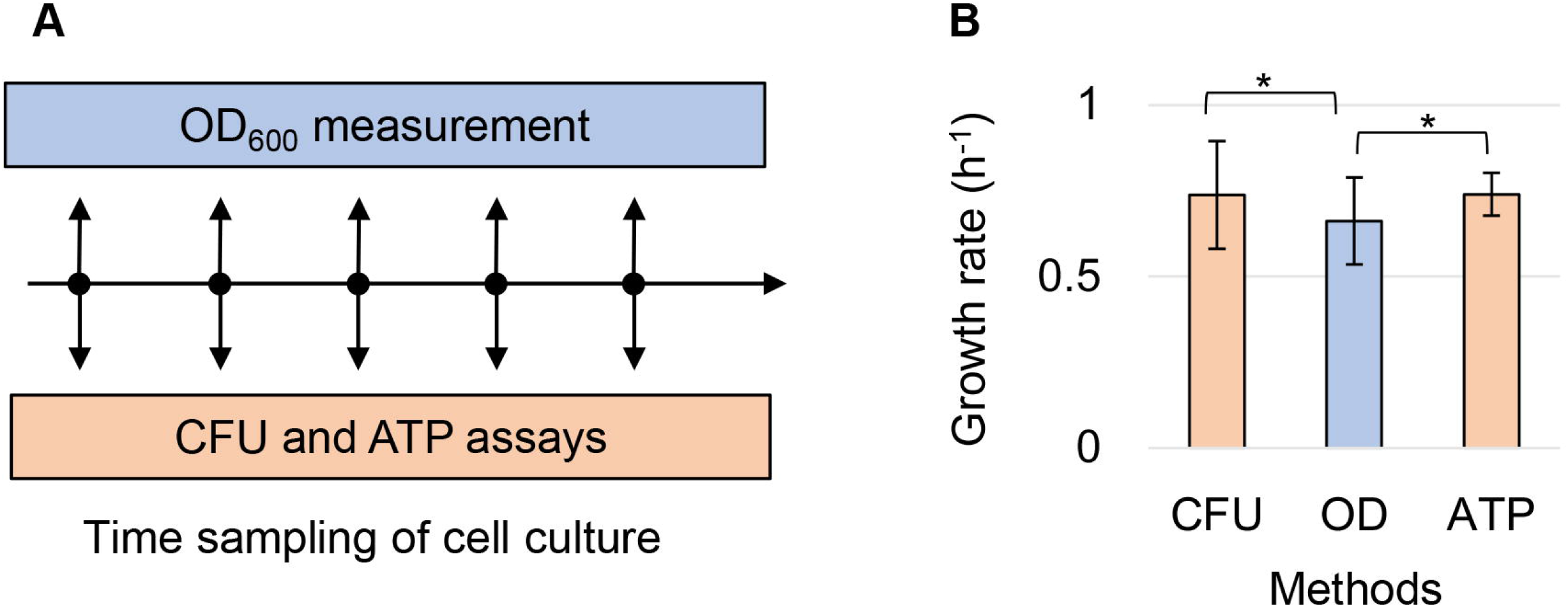
Lower estimated growth rate by means of optical turbidity. **A.** Schematic drawing of growth assays performed three different methods in parallel. Time sampling of the cell culture was performed at intervals of hours, as indicated by the arrows. The sampled cell culture was subjected to three assays based on the optical turbidity (OD_600_ measurements), colony formation (CFU) and ATP abundance (ATP). **B.** The differentiation in the growth rates evaluated by the three methods. The mean growth rates repeatedly evaluated by means of the OD_600_ (blue), CFU and ATP (orange) are shown. The standard errors of biological replication are shown. Asterisks indicate statistical significance (*P*<0.05).

### Decay effect in optical turbidity triggered the lower estimated growth rate

A decay effect was observed when the optical turbidity results were compared to the records of colony counts and OD_600_ values. Both the CFU assay (Fig. 2A, upper) and OD_600_ reads (Fig. 2A, middle) were performed in parallel to monitor the same population growth. The temporal changes in both records were supposed to be similar. Nevertheless, a temporal decrease in the ratio of the OD_600_ reads and CFU counts was observed (Fig. 2A, bottom). Since the decrease was initiated from the early exponential phase, the growth rates evaluated by optical turbidity (μ_OD_) were lower than those evaluated by colony formation (μ_CFU_) (Fig. 1B); this defense was considered to be due to the decay effect of the OD_600_ values. It was carefully tested whether the decay effect was caused by limitations of the optical measurement. No significant mechanical errors of the optical reads were detected during the exponential growth phase (OD_600_<0.5) compared with the significant decrease in the reads of OD_600_ during the stationary phase (OD_600_>1) (Fig. S1). It was verified that the decay of the OD_600_/CFU ratio during the exponential phase was not attributed to mechanical errors of the optical reads but was mainly due to biological issues, such as a difference in cellular properties.

**Figure 2.**
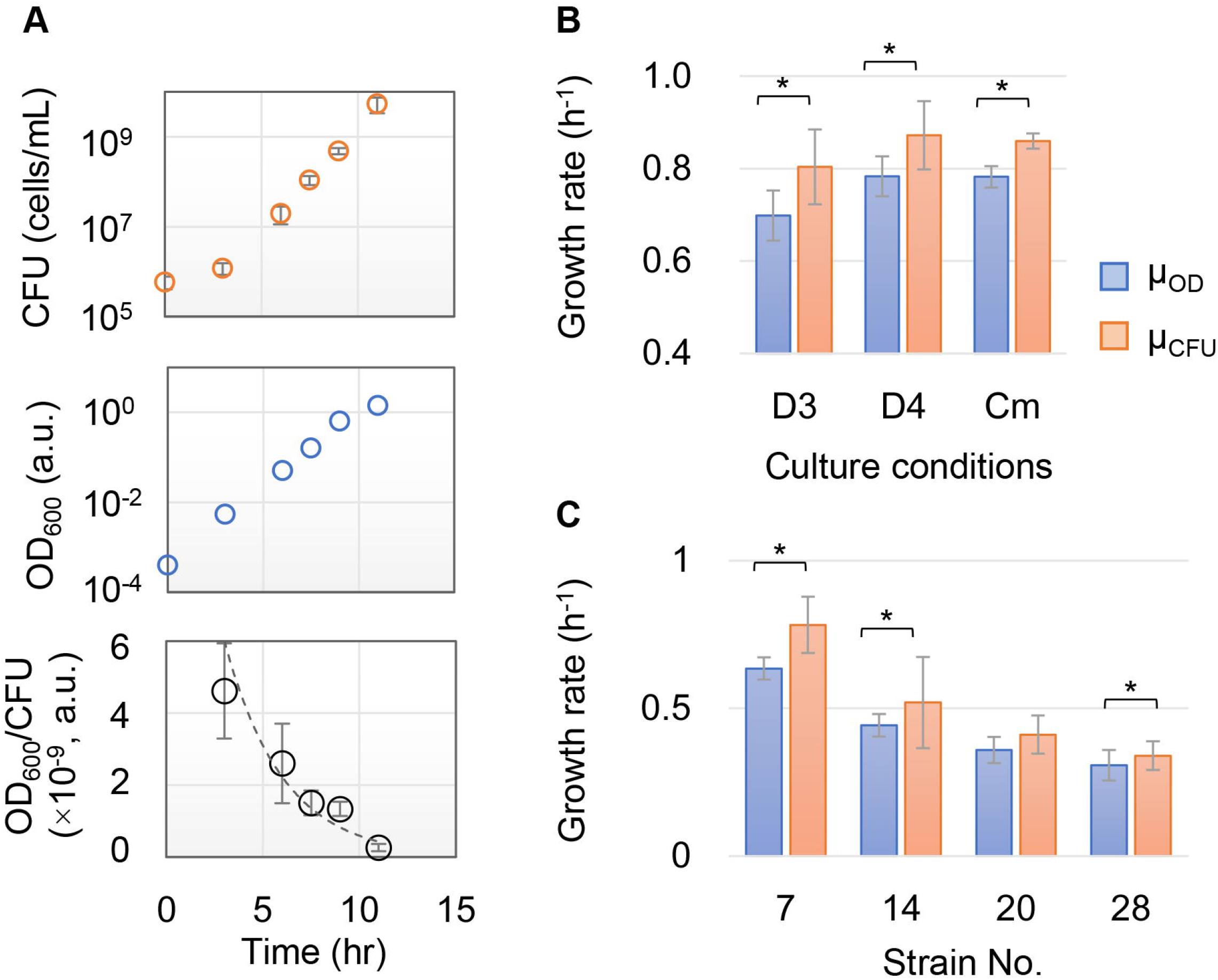
Differentiated growth rates estimated by means of the optical turbidity and colony formation. **A.** Repressed increase of OD_600_ in comparison to that of CFU. The temporal changes in CFU (upper), OD_600_ (middle), and the ratio of the two reads (bottom) of identical population growth are shown. Error bars indicate the standard errors of the assays. **B.** The growth rates under varied growth conditions. Growth of the same wild-type *E. coli* cells growing under the different conditions (D3, D4 and Cm) was analyzed in parallel by the two methods. D3 indicates that the cell growth that started from the pre-culture reached the stationary phase with a 1,000-fold dilution in fresh medium. D4 indicates that the cell growth that started from the pre-culture remained in the exponential phase with a 10,000-fold dilution in fresh medium. Cm refers to cell growth in the presence of a low concentration of antibiotics (15 μg/mL chloramphenicol). The growth rates evaluated by OD_600_ and CFU are indicated as μ_OD_ and μ_CFU_, respectively. **C.** The growth rates of the various genotypes. The growth dynamics of *E. coli* cells of four reduced genomes were analyzed by the two methods in parallel. Strain Nos. 7, 14, 20 and 28 indicate multiple deletions of approximately 251, 710, 899 and 982 kbs from the wild-type genome W3110, respectively. The standard errors of the biological replication are indicated, and asterisks indicate the statistical significance (*P*<0.05).

The lower estimated growth rate according to optical turbidity was also detected either under other growth conditions or in other genotypes. Wild-type *E. coli* cells W3110 grown under various conditions (D3, D4 and Cm described in the figure legend) also showed lower growth rates (N=3, *P*<0.05) estimated by OD_600_ (Fig. 2B). In addition, the growth dynamics of four different *E. coli* strains with reduced genomes (Strain Nos. 7, 14, 20 and 28), which were previously constructed (Mizoguchi et al., 2008) and characterized (Kurokawa et al., 2016), were assayed by the two methods (N=3~4). Similarly, the values of μ_OD_ tended to be smaller than those of μ_CFU_ (*P*<0.05) regardless of the multiple deletions of genomic sequences (Fig. 2C). The results indicated that the lower growth rates estimated by optical turbidity, which is the most commonly used method in bacterial growth studies, frequently occurred in different genotypes and under various growth conditions.

Collectively, a lower μ_OD_ than μ_CFU_ was found to be common. This result was attributed to the repressed increase in the OD_600_ value during population growth, which suggests that the changes in the cellular features were not always strictly correlated to the changes in cell number or even during the exponential growth phase. This experimental finding raised the question of whether the theoretical models that use the reads of optical turbidity properly describe bacterial growth. Because the temporal assay using OD_600_ is convenient and because it is easy to perform continuous reads in theoretical studies, a revision of the existing growth model through a consideration of the decay in OD_600_ seemed practical.

### A decay rate was introduced into the logistic model to offset the decay effect

A modified logistic model is proposed in the present study by introducing a novel parameter, the decay rate, and considering the cellular changes that accompanied growth. The logistic equation is the most popular model that is used to represent bacterial growth dynamics. This equation is commonly applied to fit growth curves based on optical turbidity, especially in systems and computational biology to understand the dynamics of living systems. Considering the decay effect in OD_600_ during population growth, the logistic model was revised. A new parameter, the decay rate (*d*), was defined, which has the same unit of h^−1^ as the growth rate.

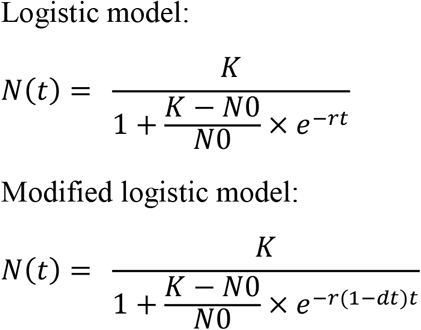

where *N*_*0*_, *K*, *r*, and *d* represent the initial cell concentration, saturated cell concentration, growth rate, and decay rate, respectively.

To evaluate whether the modified logistic model was applicable, a large number of growth data sets were fitted to the two logistic equations. A total of 710 precisely measured growth curves from 29 different *E. coli* strains in either rich (LB) or poor (M63) medium were subjected to the theoretical fitting. The histograms of parameter *K* estimated by the two models were completely overlapped (Fig. 3A). However, the histograms of the growth rate, *r*, were different between the two models (Fig. 3B). The mean growth rates estimated by the modified logistic model were larger and with a larger variation. The results indicated that the addition of the decay rate *d* in the logistic model did not influence *K*, the saturated population density or the carrying capacity, but contributed to the growth rate *r* independent of the nutritional conditions for population growth (Fig. S2).

**Figure 3.**
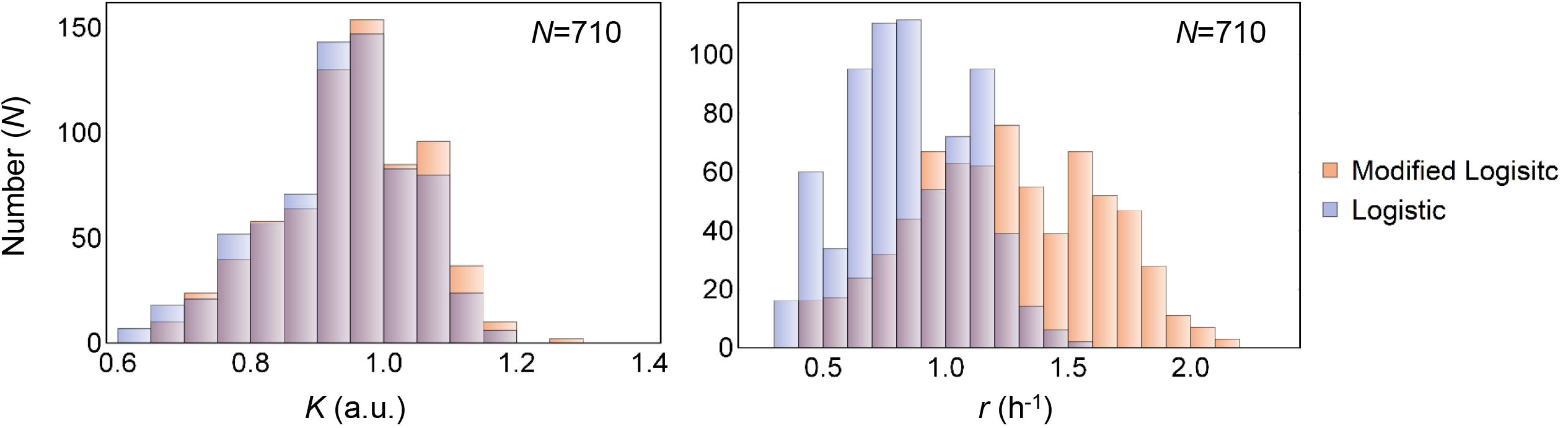
Growth parameters estimated by the two logistic models. **A.** Histograms of the saturated density *K* estimated by the two models. **B.** Histograms of the growth rate *r* estimated by the two models. Transparent blue and orange colors represent the theoretical fitting of experimental data with the logistic and modified logistic models, respectively. The number of growth curves is also indicated.

### Improved goodness of fit due to the addition of the decay rate to the logistic model

Intriguingly, the modified logistic model presented improved goodness of fit compared to the original model. The residual errors, which were designated as the sum of the squared errors (*SSE*), of fitting the 710 total growth curves to the modified logistic model were smaller than those from the original model (Fig. 4A). Thus, the goodness of fit (*R*^*2*^) was better for the modified logistic model (Fig. 4B), independent of either the genotype or medium (Fig. S3). The results showed that the modified logistic model better explained the growth dynamics of bacterial cells, which strongly indicated that the revision of the logistic model was theoretically practical. Considering the fact that the increase in OD_600_ was decided not only by exponential changes in the number of cells but also by unclear changes in the cellular contents, such as the cell size and macromolecular abundance, revision of the logistic model by introducing an additional parameter of the decay rate was also biologically reasonable.

**Figure 4.**
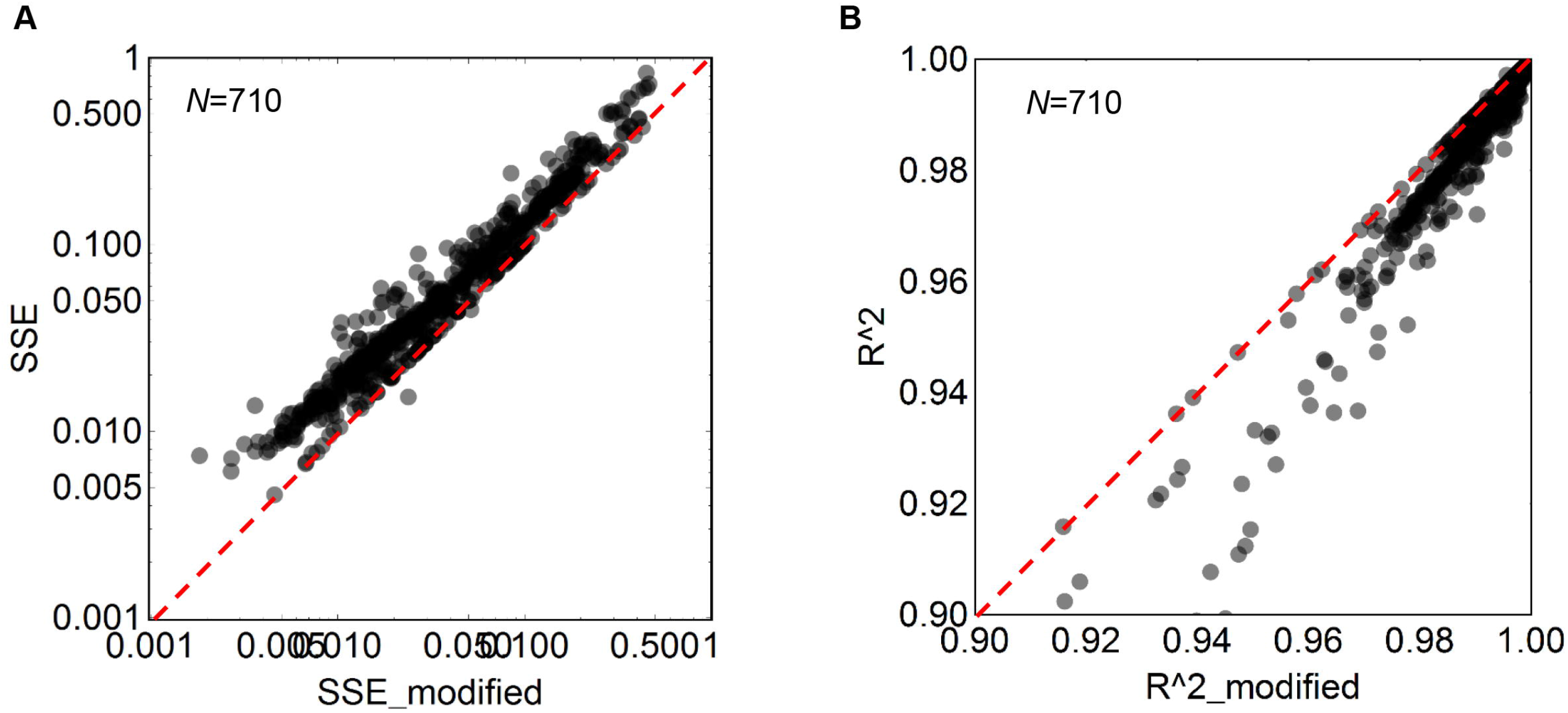
Improved fitting efficiency of the modified logistic model. **A.** A comparison of the fitting residual errors between the two models. The fitting errors are represented by the sum of squares error, *SSE*. **B.** A comparison of the goodness of fit between the two models. The goodness of fit is represented by the coefficient of determination, *R*^*2*^. The broken red lines indicate the equivalent fitting efficiency of the two models. The number of growth curves used is also indicated.

Although the parameter of *r* is known to be a constant value in the logistic model, the growth rate of μ_OD_ was somehow inconstant during the exponential phase when monitoring by the microplate reader (Fig. S4), probably due to the decay effect, which was consistent with the finding that the increase of OD_600_ was slower with population growth in comparison to that of CFU (Fig. 2A). Therefore, the parameter of *r* in the logistic equation, which presents the exponential changes of a population increase, might represent the growth rate of μ_CFU_ and changes in the number of cells without considering changes in cellular features. Because the logistic model was widely applied to growth data based on reads of OD_600_, the additional decay rate, which compensated for the decrease in μ_OD_, met the decay effect and contributed to the effective logistic fitting of bacterial growth based on optical turbidity.

### The decay rate was correlated to genome reduction in a nutritional dependent manner

Interestingly, a correlation between genome reduction and the decay rate was observed when applying the modified logistic model to the experimental bacterial growth data. Fitting the growth data of the reduced genomes to the modified logistic equation, as described above, resulted in the estimated constants of *r* and *d*, which are the growth and decay rates, respectively. Both the mean growth rates and mean decay rates of the repeated growth assays (N=12~28) were calculated for a total of 29 strains (Nos. 0~28). The results showed that a decrease along with a genome reduction was detected not only in the growth rates but also in the decay rates (Fig. 5A). The larger the genome reduction, the lower the detection of decay. This finding was supported by the fact that the changes in growth rates evaluated by OD_600_ and CFU were smaller with the genome reduction (Fig. 5B). Fewer changes between μ_OD_ and μ_CFU_ resulted in a smaller decay effect in OD_600_. Reduction of the genome size might change the cellular properties that are linked to population growth.

**Figure 5.**
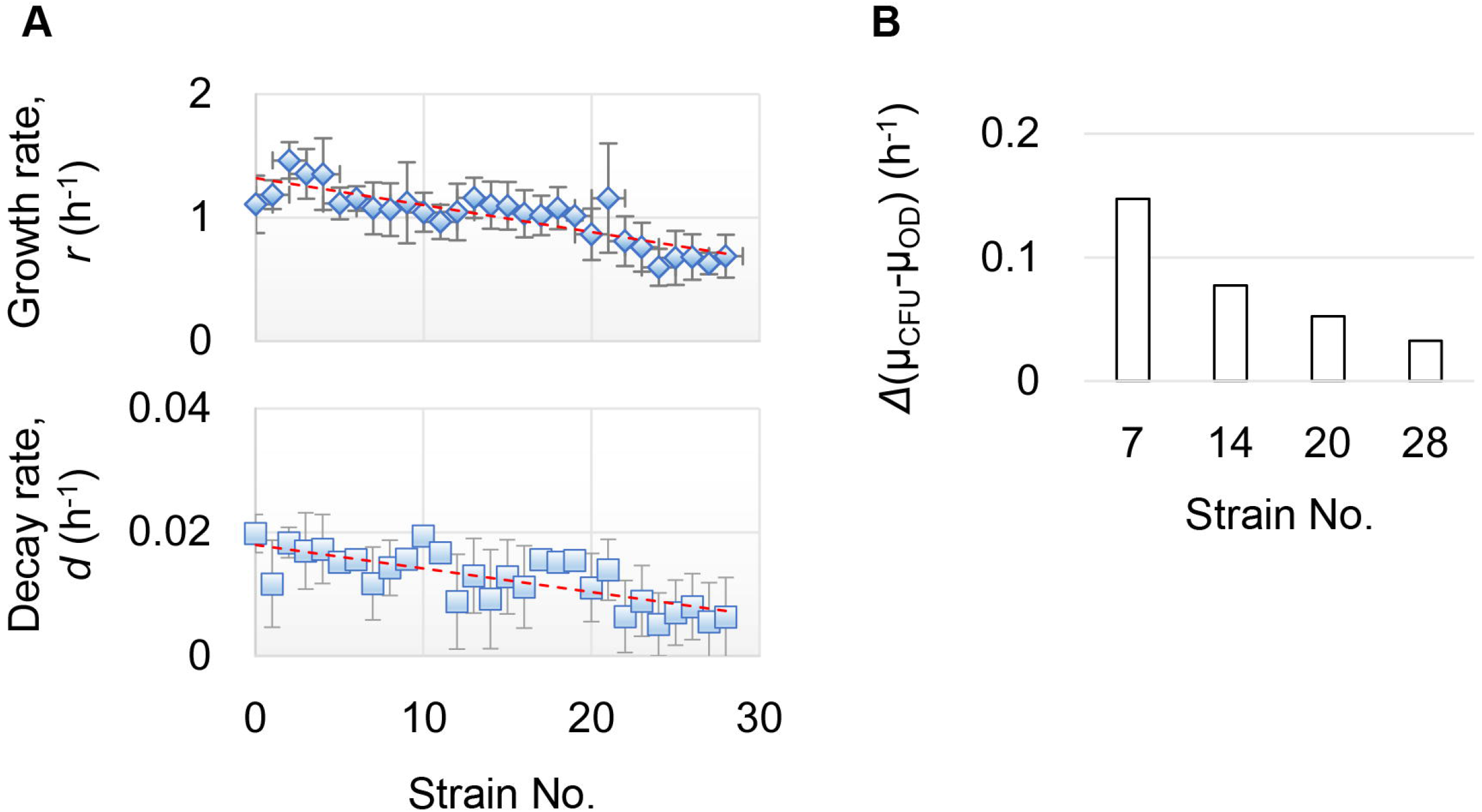
Correlation between the genome reduction and decay rate. **A.** Genome reduction dependent changes in both growth and decay rates. The mean growth rates (*r*, upper) and mean decay rates (*d*, bottom) are a result of the fitting of the growth curves in M63 with the modified logistic model and are shown based on the order of the strain No. The genomes of strain Nos. 1~28 stand for reduced genomes, and No. 0 indicates the wild-type genome W3110. The standard errors of the growth curves/tests (N=12~28) of the same strain are indicated. The broken lines in red represent the linear regression. **B.** The difference between the growth rates evaluated by CFU and OD_600_. The growth rates evaluated by OD_600_ and CFU are indicated as μ_OD_ and μ_CFU_, respectively. The changes in growth rates were calculated according to the results from Fig. 2C. The reduced genomes are shown as strain Nos. 7, 14, 20 and 28 in the order of the length of the genome reduction from short to long.

A correlation between the growth rate and genome reduction has been previously reported according to the manual calculation (Kurokawa et al., 2016). The same conclusion was drawn from the theoretical fitting suggested here, and the modified logistic model was applicable. Notably, an additional conclusion of the correlation between the decay rate and genome reduction was drawn based on the modified logistic model. The Pearson correlation coefficients of the genome size to the decay rate were highly significant (*cor*=0.684, *p*=4e-5) and were comparable to the growth rate (*cor*=0.772, *p*=6e-7). This result indicated that the changing cellular features that accompanied the population growth disturbed the optical properties. Note that the decline in the decay rates of the reduced genomes was significantly detected in the cell growth in the M63 medium but not LB medium (Fig. S5), which indicated that the nutritional richness affected the significance of the cellular changes that contributed to the decay rate. This finding agreed with the result that the difference of the fitting goodness between the two models was slight in LB but significant in M63 (Fig. S3).

### The decay rate was partially attributed to the changes in cell size

As the decay rate of the reduced genome was much smaller than that of the wild-type genome (Fig. 5), we investigated whether any biological features could be identified to explain the decrease in the decay rate. As the growth rate is often discussed with cell size (Vadia and Levin, 2015) and the genome reduction was found to somehow contribute to the change in cell size (Kurokawa et al., 2016), growth accompanying changes in cell size might be linked to the decay rate. To verify the assumption, cells carrying either the wild-type (No. 0) or the reduced (No. 28) genomes were cultured, and the size distributions at a varied population density were analyzed. Since the cell size of an identical population often showed a large variation (Fig. 6A), the median of the cell size of the population was used as a representative parameter (Yoshida et al., 2014). The results showed that the median cell size was smaller compared to the increase in population density (Fig. 6B, upper). The feature of a higher population density linked to a smaller cell size was clearly identified in the wild-type genome, but not in the reduced genome (Fig. 6B, bottom). This result suggests that the population density dependent size effect became insignificant due to genome reduction, consistent with the findings of the decay rate, and significantly declined in the reduced genome (Fig. 5). Thus, cell size changes of the growing population might be one of the reasons that cause the decay effect in OD_600_. Population density dependent changes in cell size were supposed to happen during the stationary phase due to nutritional depletion (Vadia and Levin, 2015; Ying et al., 2015); however, the results showed that the changes in cell size also occurred even in the exponential phase. This result agrees well with the temporal decrease in the ratio of OD_600_/CFU from the exponential phase (Fig. 2A).

**Figure 6.**
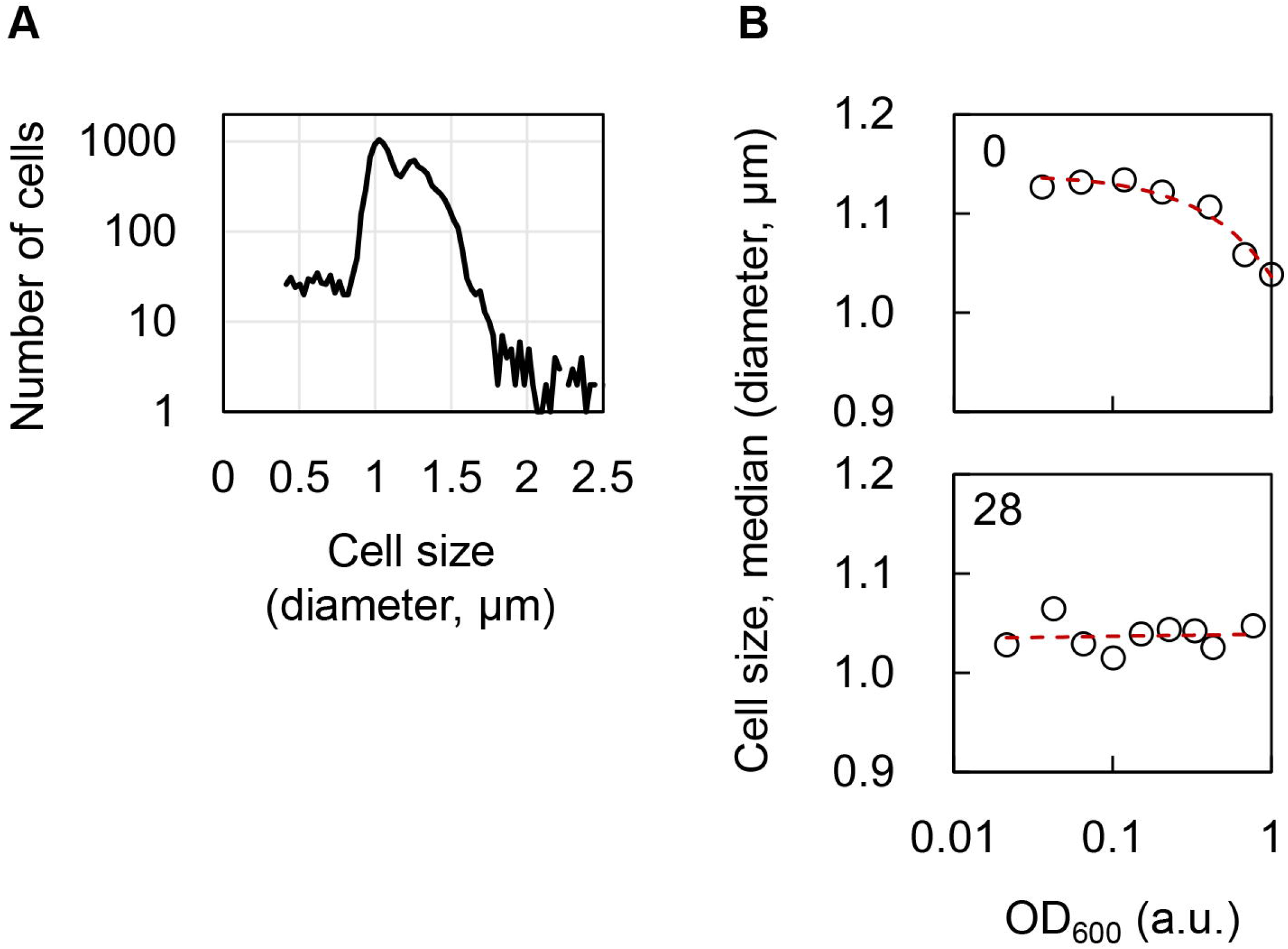
Relationship between the cell size and population density. **A.** Size distribution of an identical cell population. A size distribution of more than 10,000 *E. coli* cells growing exponentially in M63 is shown as an example. **B.** The relationship between the population density and cell size. *E. coli* cells of either the wild-type (No. 0, upper) or the reduced (No. 28, bottom) genomes were grown in M63. The cell size of the culture at various population densities was measured, and the median size of the cell population was used for analysis. The red broken lines represent the exponential regression.

The optical turbidity might reflect both the cell number and cell size. The decay rate (*i.e.*, decay effect in optical turbidity) could be partially explained by changes in the cell size/volume accompanied by the population increase, and the significance of the cell size-mediated decay effect was genome/strain dependent. The modified logistic model included an additional parameter considering the cellular features, such as the cell size, to compensate for the original logistic model, which is often applied to population growth with the quantitative unit of the cell number.

In summary, a decay effect in the bacterial growth rate evaluated by optical turbidity was detected according to the differentiation in growth rates caused by variations in methods. To compensate for the decay effect in OD_600_ reads, the commonly used growth model of the logistic equation was revised by introducing a new parameter, the decay rate, which was found to be dependent on both the genotype and medium and was partially attributed to changes in cell size. These new findings reveal the biological impact of considering the decay effect in the bacterial growth rate evaluated by optical turbidity. The novel finding of the correlation of genome reduction to both the growth and the decay rates is highly intriguing, which requiring further

## Materials and Methods

### Strains and media

The *Escherichia coli* strain W3110 and its derivative strains (Mizoguchi et al., 2008; Kurokawa et al., 2016) used in the present study were acquired from the National BioResource Project (NBRP) of the National Institute of Genetics (NIG), Japan. *Escherichia coli* cells were cultured in minimal medium M63, which contains 62 mM dipotassium hydrogenphosphate, 39 mM potassium dihydrogen phosphate, 15 mM ammonium sulfate, 15 μM thiamine hydrochloride, 1.8 μM Iron (II) sulfate, 0.2 mM magnesium sulfate, and 22 mM glucose. Preparation of the M63 medium was previously described in detail (Kurokawa and Ying, 2017). The colony formation was detected using LB broth agar, Miller (Sigma Aldrich).

### Bacterial growth in the test tubes

A total of 43 mL of M63 medium was placed in a 50 ml Falcon (Watson) tube in which 100 μL of the *Escherichia coli* glycerol stock stored at −80 °C was added and well mixed using a vortex. A total of 5 mL of the mixture was dispensed into eight sterilized glass tubes and cultured at 37 °C and 200 rpm using a bioshaker (Taitec). The culture was temporally sampled from the late lag phase to the early stationary phase to detect the growing population using three different methods that relied on the physical, biological and chemical properties of bacterial cells, *i.e.*, the optical turbidity, colony formation, and ATP abundance. Sampling was conducted by taking 1.2 mL of the culture from two out of eight glass tubes, and 1 mL of the sampled culture was placed in a cuvette to measure absorbance at OD_600_ (Beckman DU730), and the remaining volume was used for the other two assays. Time sampling was performed rotationally among a total of eight glass tubes, which led to a single growth curve based on the OD_600_ value. This time course of bacterial growth was conducted repeatedly (N=3~5). The biological replications varied from the initial pre-culture.

### Colony forming unit (CFU) assay

The colony formation (colony forming unit, CFU) assay is a method of estimating the number of viable bacteria in a culture solution by counting colonies on the assumption that a microorganism colony that appears on agar medium is derived from one cell. According to the value of OD_600_, multiple dilution rates of the sampled culture, which varied in order, were performed to obtain 10 to 500 colonies per plate on which 100 μL of the diluted culture was plated. The plates were cultured overnight in an incubator (Yamato) at 37 °C, and then, a colony count was performed. At least four plates were used for each sampling point, and approximately 40 plates were used for a single growth curve based on the CFU. Three to five biological replications of the growth curves were performed for each strain or culture condition, which resulted in the consumption of approximately 2,000 LB agar plates in the present study.

### Measurement of the cellular ATP concentration

Six different concentrations of the ATP solutions were prepared for the standard curve by diluting 10.4 mg/mL ATP (Roche) 10^4^~10^9^-fold with M63 medium. After placing 100 μL of M63 medium or the six ATP solutions in 1.5-mL microtubes, 100 μL of BacTiter-Glo ™ Microbial Cell Viability Assay solution (Promega) was added to each microtube according to the manufacturer’s instructions. The fluorescence intensity was measured to prepare a calibration curve. A total of 10, 20, 50, and 100 μL of each sampled culture was measured similarly to estimate the amount of ATP in the growing cells. Based on the calibration curve, the cellular ATP concentration was determined. A growth curve based on the ATP concentration was made to calculate the growth rate.

### Calculation of the growth rate

Growth curves based on three different methods were created according to the temporal measurements of OD_600_, CFU, and/or ATP as described above. A simple exponential regression toward the exponential growth phase in the growth curve was performed as follows.

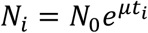

where *N*, *t* and *μ* represent the population density (cell concentration), time (h) and growth rate, respectively. Regression resulted in an exponent of the exponential slope as the growth rate (h^−1^). The growth rates calculated from the growth curves of OD_600_, CFU and ATP were named μ_OD_, μ_CFU_ and μ_ATP_, respectively.

### Detection of the cell size

The cell culture (glycerol stock) was repeatedly diluted twofold with M63 medium, which led to a series of dilution rates from two to 1,028-fold. A total of 2 mL of the diluted cell mixtures was loaded on a 24-well microplate (Iwaki). The microplate was incubated for 12 h in a microplate mixer (Taitec) with a rotation rate of 500 rpm at 37 °C as described previously (Nishimura et al., 2017). All of the wells (cell cultures) were subjected to measurements by both optical turbidity and a cell counter to reach a correlation between the OD600 and the mean cell size of the cell population. OD600 was measured using a 1 mL cuvette as described above. The cell size (volume) of the growing *E. coli* cells were measured using a cell counter (Beckman, Multisizer 4) as described previously (Kurokawa et al., 2016).

### Bacterial growth in a 96-well microplate

Glycerol stocks of the *E. coli* cultures collected at the early exponential phase (OD_600_<0.05) were diluted 100- to 1000-fold in M63 medium and well mixed using a vortex. The diluted cell mixture was loaded onto a 96-well microplate (Costar) in multiple wells at varied locations with 200 μL of mixture per well as previously described (Kurokawa and Ying, 2017). The 96-well microplate was incubated in a plate reader (Epoch2, BioTek) with a rotation rate of 600 rpm at 37 °C. Bacterial growth was detected at an absorbance of 600 nm and read at intervals of 5 min. The growth rates were calculated from every two reading points in a growth curve and according to the following equation.

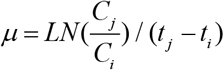

where *C*_*i*_ and *C*_*j*_ represent the two reads of OD_600_ values at two time points of *t*_*j*_ and *t*_*i*_, respectively, which were discontinuous at an interval of 1 h (*i.e.*, every 20 intervals), although the reads were at intervals of 5 min.

### Computational analysis of the bacterial growth

The theoretical models for bacterial growth were evaluated using Python. The logistic and modified logistic models were fitted with 710 growth curves of 29 *Escherichia coli* strains of various genome sizes growing in 96-well microplates. The raw data sets of these growth curves can be found at Springer Nature Research Data Support and can be accessed online *via* the following URL: https://doi.org/10.6084/m9.figshare.5918608. The details regarding the determination of the growth data set was previously described (Kurokawa et al., 2016; Kurokawa and Ying, 2017), and the data sets of growth in M63 and LB media were used in the present study. Evaluation of the models (goodness of fit) was assessed based on the sum of squares, which were calculated according to the following equations.

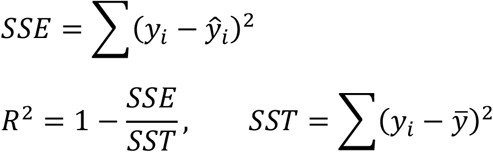

where *SSE*, *SST* and *R*^*2*^ represent the sum of squares error, the sum of squares total, and the coefficient of determination, respectively.

## Acknowledgments

This study was partially supported by the Tsukuba Basic Research Support Program (Type B). The funder had no role in the study design, data collection and interpretation or the decision to submit the work for publication.

## Author Contributions

KT, MK and HA performed the experiments; YYC, YT and BWY performed the analyses; BWY conceived the research and wrote the paper. All authors read and proved the final manuscript.

## References

Alonso, A.A., Molina, I., and Theodoropoulos, C. (2014). Modeling bacterial population growth from stochastic single-cell dynamics. Appl Environ Microbiol 80(17), 5241–5253. doi: 10.1128/AEM.01423-14.

Dalgaard, P., Ross, T., Kamperman, L., Neumeyer, K., and McMeekin, T.A. (1994). Estimation of bacterial growth rates from turbidimetric and viable count data. Int J Food Microbiol 23(3-4), 391–404.

Desmond-Le Quemener, E., and Bouchez, T. (2014). A thermodynamic theory of microbial growth. Isme j 8(8), 1747–1751. doi: 10.1038/ismej.2014.7.

Egli T. (2015). Microbial growth and physiology: a call for better craftsmanship. Front Microbiol 6, 287. doi: 10.3389/fmicb.2015.00287.

Fujikawa, H., and Morozumi, S. (2005). Modeling surface growth of Escherichia coli on agar plates. Appl Environ Microbiol 71(12), 7920–7926. doi: 10.1128/AEM.71.12.7920-7926.2005.

Hall, B.G., Acar, H., Nandipati, A., and Barlow, M. (2014). Growth rates made easy. Mol Biol Evol 31(1), 232–238. doi: 10.1093/molbev/mst187.

Harris, C.M., and Kell, D.B. (1985). The estimation of microbial biomass. Biosensors 1(1), 17–84.

Hermsen, R., Okano, H., You, C., Werner, N., and Hwa, T. (2015). A growth-rate composition formula for the growth of E.coli on co-utilized carbon substrates. Mol Syst Biol 11(4), 801. doi: 10.15252/msb.20145537.

Kargi F. (2009). Re-interpretation of the logistic equation for batch microbial growth in relation to Monod kinetics. Lett Appl Microbiol 48(4), 398–401. doi: 10.1111/j.1472-765X.2008.02537.x.

Koseki, S., and Nonaka, J. (2012). Alternative approach to modeling bacterial lag time, using logistic regression as a function of time, temperature, pH, and sodium chloride concentration. Appl Environ Microbiol 78(17), 6103–6112. doi: 10.1128/AEM.01245-12.

Kurokawa, M., Seno, S., Matsuda, H., and Ying, B.W. (2016). Correlation between genome reduction and bacterial growth. DNA Res 23(6), 517–525. doi: 10.1093/dnares/dsw035.

Kurokawa, M., and Ying, B.W. (2017). Precise, High-throughput Analysis of Bacterial Growth. J Vis Exp (127). doi: 10.3791/56197.

Lin, H.L., Lin, C.C., Lin, Y.J., Lin, H.C., Shih, C.M., Chen, C.R., et al. (2010). Revisiting with a relative-density calibration approach the determination of growth rates of microorganisms by use of optical density data from liquid cultures. Appl Environ Microbiol 76(5), 1683–1685. doi: 10.1128/AEM.00824-09.

Liu, B., Liu, H., Pan, Y., Xie, J., and Zhao, Y. (2016). Comparison of the Effects of Environmental Parameters on the Growth Variability of Vibrio parahaemolyticus Coupled with Strain Sources and Genotypes Analyses. Front Microbiol 7, 994. doi: 10.3389/fmicb.2016.00994.

Madrid, R.E., and Felice, C.J. (2005). Microbial biomass estimation. Crit Rev Biotechnol 25(3), 97–112. doi: 10.1080/07388550500248563.

Mizoguchi, H., Sawano, Y., Kato, J., and Mori, H. (2008). Superpositioning of deletions promotes growth of Escherichia coli with a reduced genome. DNA Res 15(5), 277–284. doi: 10.1093/dnares/dsn019.

Myers, J.A., Curtis, B.S., and Curtis, W.R. (2013). Improving accuracy of cell and chromophore concentration measurements using optical density. BMC Biophys 6(1), 4. doi: 10.1186/2046-1682-6-4.

Nishimura, I., Kurokawa, M., Liu, L., and Ying, B.W. (2017). Coordinated Changes in Mutation and Growth Rates Induced by Genome Reduction. MBio 8(4). doi: 10.1128/mBio.00676-17.

Peleg, M., and Corradini, M.G. (2011). Microbial growth curves: what the models tell us and what they cannot. Crit Rev Food Sci Nutr 51(10), 917–945. doi: 10.1080/10408398.2011.570463.

Pla, M.L., Oltra, S., Esteban, M.D., Andreu, S., and Palop, A. (2015). Comparison of Primary Models to Predict Microbial Growth by the Plate Count and Absorbance Methods. Biomed Res Int 2015, 365025. doi: 10.1155/2015/365025.

Ponciano, J.M., Vandecasteele, F.P., Hess, T.F., Forney, L.J., Crawford, R.L., and Joyce, P. (2005). Use of stochastic models to assess the effect of environmental factors on microbial growth. Appl Environ Microbiol 71(5), 2355–2364. doi: 10.1128/AEM.71.5.2355-2364.2005.

Roessler, M., Sewald, X., and Muller, V. (2003). Chloride dependence of growth in bacteria. FEMS Microbiol Lett 225(1), 161–165.

Sprouffske, K., and Wagner, A. (2016). Growthcurver: an R package for obtaining interpretable metrics from microbial growth curves. BMC Bioinformatics 17, 172. doi: 10.1186/s12859-016-1016-7.

Swain, P.S., Stevenson, K., Leary, A., Montano-Gutierrez, L.F., Clark, I.B., Vogel, J., et al. (2016). Inferring time derivatives including cell growth rates using Gaussian processes. Nat Commun 7, 13766. doi: 10.1038/ncomms13766.

Tonner, P.D., Darnell, C.L., Engelhardt, B.E., and Schmid, A.K. (2017). Detecting differential growth of microbial populations with Gaussian process regression. Genome Res 27(2), 320–333. doi: 10.1101/gr.210286.116.

Vadia, S., and Levin, P.A. (2015). Growth rate and cell size: a re-examination of the growth law. Curr Opin Microbiol 24, 96–103. doi: 10.1016/j.mib.2015.01.011.

Verhulst P.F. (1845). Recherches mathématiques sur la loi d’accroissement de la population. Nouv. mém. de l’Academie Royale des Sci. et Belles-Lettres de Bruxelles 18, 1–41.

Verhulst P.F. (1847). Deuxième mémoire sur la loi d’accroissement de la population. Mém. de l’Academie Royale des Sci., des Lettres et des Beaux-Arts de Belgique 20, 1–32.

Verissimo, A., Paixao, L., Neves, A.R., and Vinga, S. (2013). BGFit: management and automated fitting of biological growth curves. BMC Bioinformatics 14, 283. doi: 10.1186/1471-2105-14-283.

Winsor C.P. (1932). The Gompertz Curve as a Growth Curve. Proc Natl Acad Sci U S A 18(1), 1–8.

Yates, G.T., and Smotzer, T. (2007). On the lag phase and initial decline of microbial growth curves. J Theor Biol 244(3), 511–517. doi: 10.1016/j.jtbi.2006.08.017.

Ying, B.W., Honda, T., Tsuru, S., Seno, S., Matsuda, H., Kazuta, Y., et al. (2015). Evolutionary Consequence of a Trade-Off between Growth and Maintenance along with Ribosomal Damages. PLoS One 10(8), e0135639. doi: 10.1371/journal.pone.0135639.

Yoshida, M., Tsuru, S., Hirata, N., Seno, S., Matsuda, H., Ying, B.W., et al. (2014). Directed evolution of cell size in Escherichia coli. BMC Evol Biol 14(1), 257. doi: 10.1186/s12862-014-0257-1.

Ziv, N., Siegal, M.L., and Gresham, D. (2013). Genetic and nongenetic determinants of cell growth variation assessed by high-throughput microscopy. Mol Biol Evol 30(12), 2568–2578. doi: 10.1093/molbev/mst138.

